# Semi-automated computational assessment of cancer organoid viability using rapid live-cell microscopy

**DOI:** 10.1101/2021.12.07.471003

**Authors:** Joseph D. Buehler, Cylaina E. Bird, Milan R. Savani, Lauren C. Gattie, William H. Hicks, Michael M. Levitt, Mohamad El Shami, Kimmo J. Hatanpaa, Timothy E. Richardson, Samuel K. McBrayer, Kalil G. Abdullah

## Abstract

The creation of patient-derived cancer organoids represents a key advance in preclinical modeling and has recently been applied to a variety of human solid tumor types. However, conventional methods used to assess cellular viability in tissue specimens are poorly suited for the evaluation of cancer organoids because they are time-intensive and involve tissue destruction. To address this issue, we established a suite of 3-dimensional patient-derived glioma organoids, treated them with chemoradiotherapy, stained organoids with non-toxic cell dyes, and imaged them using a rapid laser scanning confocal microscopy method termed “Apex Imaging”. We then developed and tested a fragmentation algorithm to quantify heterogeneity in the topography of the organoids as a potential surrogate marker of viability. This algorithm, SSDquant, provides a 3-dimensional visual representation of the organoid surface and a numerical measurement of the sum-squared distance (SSD) from the derived mass center of the organoid. We tested whether SSD scores correlate with traditional immunohistochemistry-derived cell viability markers (cellularity and cleaved caspase 3 expression) and observed statistically significant associations between them using linear regression analysis. Our work describes a quantitative, non-invasive approach for the serial measurement of patient-derived cancer organoid viability, thus opening new avenues for the application of these models to studies of cancer biology and therapy.

## Introduction

Patient-derived cancer organoids are 3-dimensional preclinical models that are derived from tumor resection surgeries and recapitulate characteristics of parental tumors^1,2^. Successful generation of organoids from primary tissue samples represents a significant advance in the ability to accurately model human solid tumors in the laboratory^3,4^. However, due to the physical and molecular complexity of these organoids, using these models to test the efficacy of experimental therapies is currently limited by the need for tissue fixation, immunohistochemical staining, and subsequent quantitation^5,6^. These constraints prevent the routine surveillance of organoid viability prior to experimentation as well as longitudinal assessments of treatment response. Therefore, the development of new methods for non-invasive, serial organoid viability quantification would create new opportunities to use these cultures to test new anticancer therapies and enhance the reproducibility of data generated from these models.

Gliomas are the most common malignant brain tumors and have been particularly challenging to accurately model in the laboratory due to their cellular heterogeneity and complexity. Recent work has shown that glioblastoma organoids can be cultured directly from the operating room as live tissue that can be propagated in vitro for extended periods of time ^1,7^. Based on visual observations of patient-derived glioma organoids undergoing therapy-induced cell killing, we hypothesized that changes in organoid surface topography could serve as an alternative and readily quantifiable biomarker of the viability of organoid-resident tumor cells. To test this hypothesis, we created a rapid microscopy protocol and companion algorithm to measure topographical heterogeneity in patient-derived organoids undergoing treatment with radiation and/or anticancer drug therapies.

## Materials and Methods

### Human Subjects

Patient tissue and blood were collected following ethical and technical guidelines on the use of human samples for biomedical research at University of Texas Southwestern Medical Center after informed patient consent under a protocol approved by the University of Texas Southwestern Medical Center’s Institutional Review Board. All patient samples were de-identified before processing.

### Organoid Culture and Immunohistochemistry

Organoids were established and propagated using previously described methods with a modification to oxygen concentration within the organoid tissue incubator as described^7^. Glioma organoids were kept in 5% O_2_ and evaluated over 7 days in four cohorts: DMSO-treated (control), radiation, temozolomide or Olaparib (chemotherapeutics), and combined chemotherapy and radiation. Organoids were irradiated at a total dose of 10 Gray (Gy) at a dose rate of 2.45 Gy/min using a XRAD320 cabinet irradiator with 250 kV voltage, 15 mA current, and 65 source to surface distance. Organoids were cultured for 7 days in media containing either 50 uM temozolomide (SigmaAldrich), 5 uM Olaparib (Selleck) or DMSO. Media containing fresh drug was replaced every 48 hours. After treatment and imaging, organoids were fixed in 10% formalin for 1 hour then washed and suspended in 70% ethanol. Histology was performed by HistoWiz Inc. (histowiz.com). Samples were processed, embedded in paraffin, and sectioned at 4 μm. Immunohistochemistry was performed on a Bond Rx autostainer (Leica Biosystems). Slides were stained with an anti-cleaved caspase 3 antibody (Cell□Signaling□Technologies CST9661 1:300) and counterstained with hematoxylin to detect cell nuclei. Chromagen development was done using the Bond Polymer Refine Detection Kit (Leica Biosystems) according to the manufacturer’s protocol. After staining, sections were dehydrated and film coverslipped using a TissueTek-Prisma Coverslipper. Whole slide scanning (40x) was performed on an Aperio AT2 (Leica Biosystems). IHC quantification was performed using QuPath software (version 0.2.3). Scanned brightfield images of immunohistochemistry slides were analyzed as svs files in the opensource software package QuPath^8^. The WatershedCellDetection plugin was run and the periphery was defined by running the DilateAnnotationPlugin with a radius of -200.0 μm. Cellularity (hematoxylin+ objects/μm^2^) and cell death rate (% cleaved caspase 3-positive cells) metrics were quantified in the periphery of organoids.

### Apex Imaging

Hoescht□33342 (360/460 nm) and propidium iodide (535/617 nm) (Ready Probes Kit,□ThermoFisher□R37610) dyes were added according to manufacturer protocol and incubated for 5 minutes in the dark. Fluorescent confocal images were captured on a Zeiss LSM 780 inverted microscope at 10X magnification. Images were deconvolved with AutoQuant X3 using a blind point spread function with high noise settings and 10 iterations. An executable application was created to semi-automate the following analysis steps: deconvolved Z stacks were loaded into ImageJ, duplicated, and one copy filtered with a Gaussian Blur (Σ=2). Then the blurred stack was subjected to Classic Watershed Segmentation^9^ from the MorphoJLib to define borders between objects^10^.

### SSDquant Algorithm Development

To ensure efficient separation of overlapping nuclei, we added boundaries between local intensity maxima using a 3D Watershed algorithm. Next, we optimized the 3D Object Counter tool in ImageJ to detect each individual nucleus in 3D space. The tool was optimized by statistical determination of the intensity threshold needed to count the nuclei. The 3D Object counter exports a series of data including the volume, surface area, geometric center and intensity weighted center of each object. The intensity-weighted center coordinate matrix was selected for further analysis because it best represented the cell center of mass. This matrix of coordinates was then brought into MATLAB for surface analysis. The MATLAB curve fitting toolbox was used to optimize surface fitting to the centers of mass data. The thin plate spline approach was selected to accommodate the high data point output and render a function suitable for roughness calculations. The thin plate spline fitting was automated with the MATLAB function *tpaps. tpaps* works by optimizing the objective function:

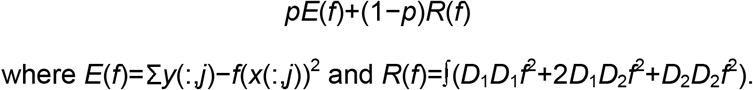

The *p* value allows the function to adapt to surface characteristics while minimizing error. A *p* value of was chosen based on the mean of the automatically assigned p value from derived images. Once the *p* value was established, we calculated the deviation as a measure of roughness. This deviation represented the sum squared distance (SSD) from each nucleus to the best fit spline generated by *tpaps*. Therefore, the SSD metric reflects the topographical heterogeneity of organoid surfaces.

### Statistics

Linear regression analyses were used to quantify correlations between natural logarithm-transformed SSD values and 1) cellularity values or 2) cell death rates in a collection of 14 organoids. Reported *p* values are derived from statistical testing of the null hypothesis that slopes are equal to zero. *p* values less than 0.05 were considered to be significant.

## Results

We created a suite of patient-derived glioma organoids from two primary tumor tissue specimens collected at the University of Texas Southwestern Medical Center. Following organoid establishment, propagation, and treatment, organoids were either fixed for immunohistochemistry (IHC) analysis and/or subjected to live-cell imaging via laser scanning confocal microscopy (Figure 1).

**Figure 1.**
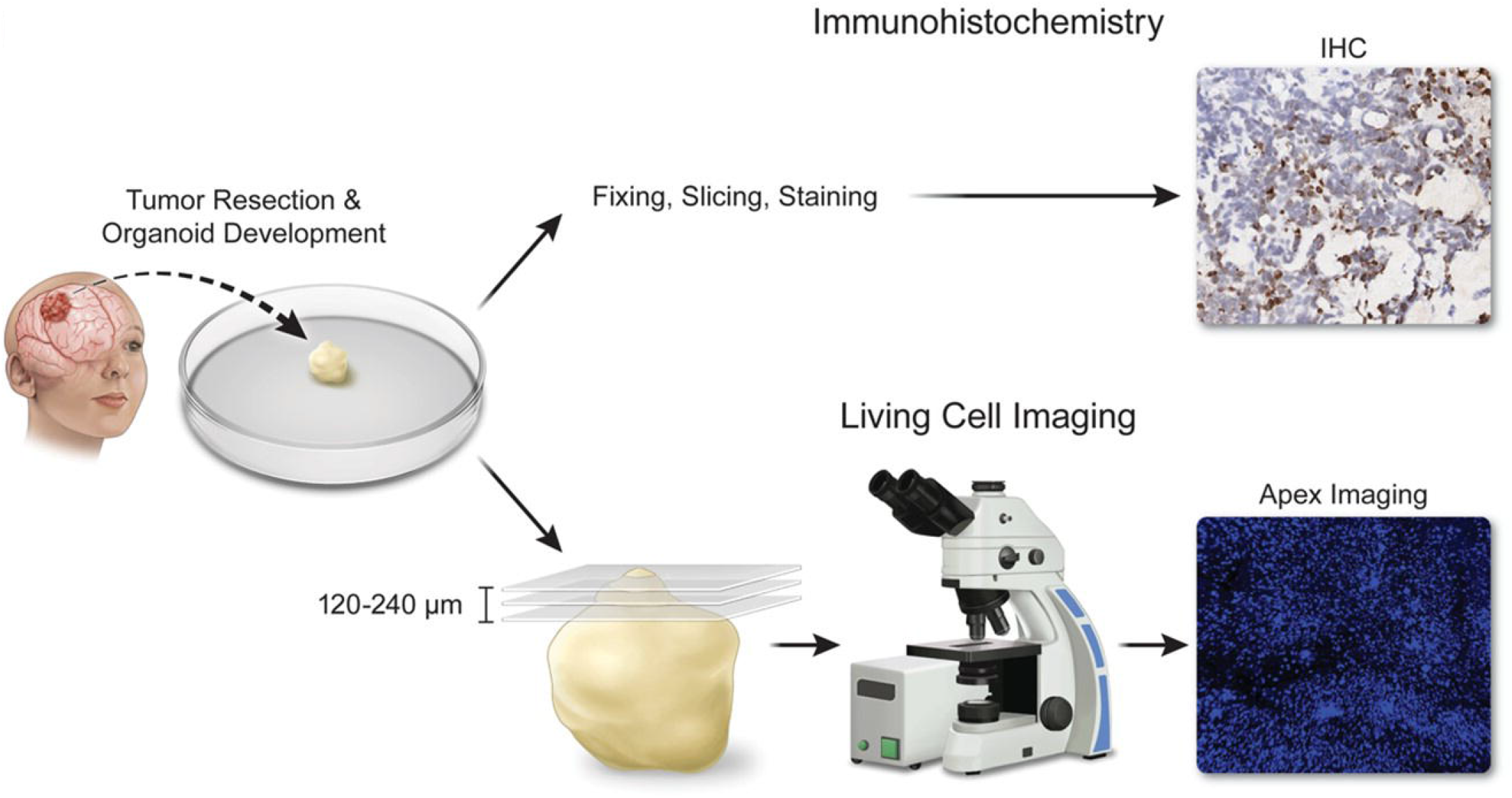
Schema of organoid comparison and analysis. Glioblastoma organoids were derived from craniotomies for brain tumors and propagated in vitro. Organoids were treated with chemoradiotherapy prior to organoid viability assessments via Apex Imaging live cell confocal microscopy and/or immunohistochemistry. Immunohistochemical analysis results were then compared head-to-head with results from tandem Apex Imaging and application of the SSDquant algorithm.

To begin developing a non-invasive method for organoid viability quantification, we conducted preliminary studies using a glioblastoma (IDH WT) organoid model. We elected to use non-toxic, fluorogenic Hoescht□33342 and propidium iodide dyes that reversibly stain live and dead cells, respectively. Importantly, discrimination of live and dead cells with this combination of dyes does not require tissue fixation or cell membrane permeabilization, thereby obviating tissue destruction during organoid viability assessment. We first stained an untreated glioblastoma organoid with Hoescht□33342 and propidium iodide dyes. Next, we performed laser scanning confocal microscopy to collect images at specific planes (Z stacks) 120-240 μm below the apex of the organoid (Figure 2A). This methodology, which we call Apex Imaging, enables precise assessment of organoid topography. Importantly, the vast majority of cells in the organoid region assayed stained positive for the live-cell Hoescht 33342 dye (shown in blue) and excluded the dead-cell propidium iodide dye (shown in red), thereby supporting the feasibility of our approach.

**Figure 2.**
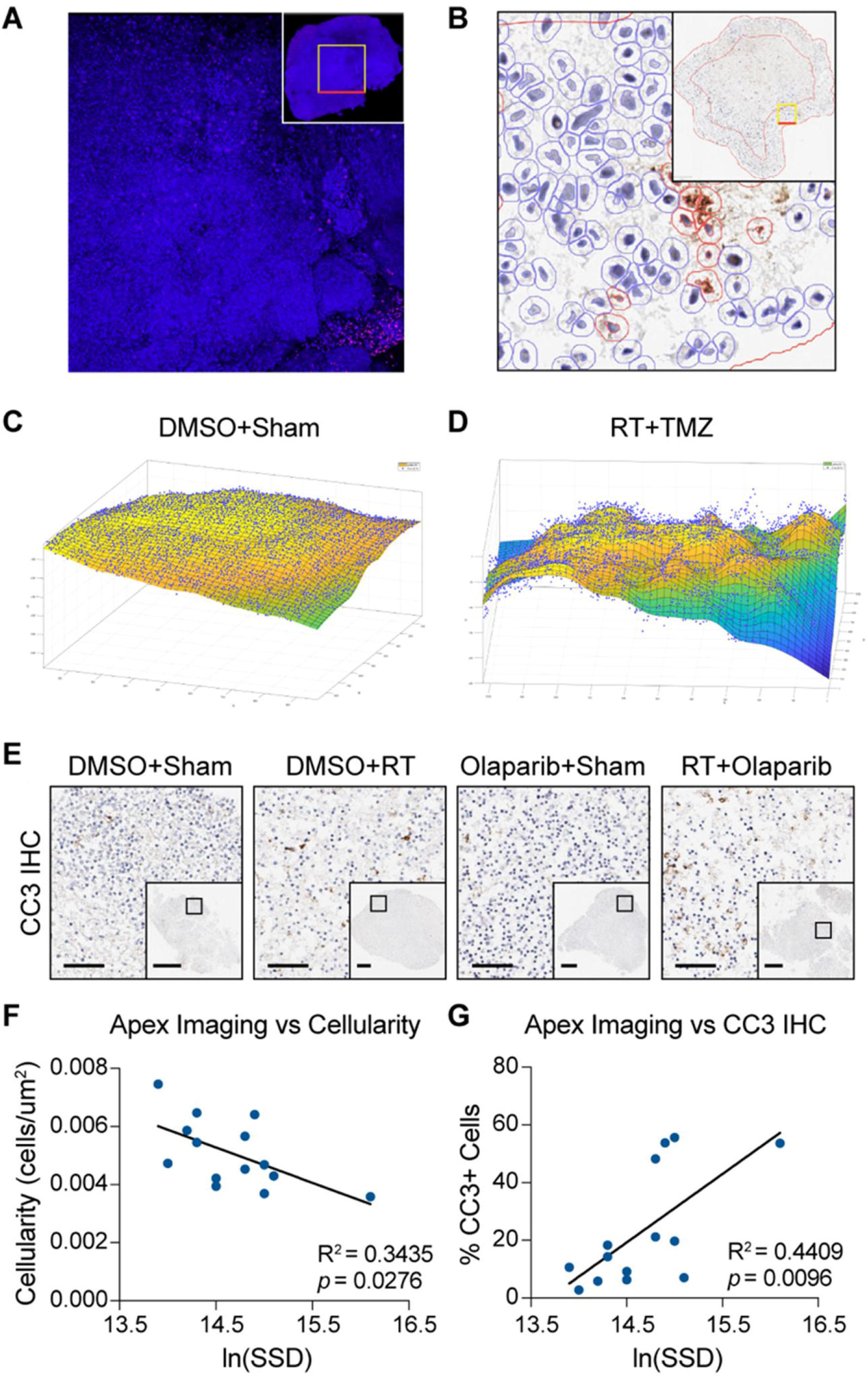
Applying the SSDquant algorithm to Apex Imaging data provides a metric of cancer organoid viability that correlates with conventional viability measurements. (**A**) Apex Imaging of untreated glioblastoma organoids stained with Hoescht 33342 and propidium iodide dyes. Live cells and dead cells are represented by blue and red puncta, respectively. (**B**) A glioblastoma organoid was irradiated, fixed, and stained with hematoxylin and an anti-CC3 antibody to evaluate cellular viability. Identification of live (blue) and dead (red) cells was performed using QuPath software. In A and B, red scale bar in insets = 200μm. Apex imaging data from untreated (**C**) or RT/TMZ dual-treated (**D**) glioblastoma organoids was processed using the SSDquant algorithm. Visual representations of organoid topography are shown. Surface color indicates the natural logarithm of SSD values in a 20×20μm region. Blue dots represent cell centers of mass. (**E**) Anaplastic oligodendroglioma organoids were treated with Olaparib, RT, both, or neither and CC3 IHC was performed to quantify cellularity and cell death rates in organoid cultures. Scale bars in insets and main figures are 1 mm and 100 μm, respectively. Linear regression analyses of organoid SSD scores versus (**F**) cellularity values or (**G**) cell death rates were conducted. Analyses revealed statistically significant correlations between SSD scores and decreasing cellularity and increasing cell death rates. In A, B, and E, boxes in insets are displayed at high magnification in main figures.

We next fixed untreated glioblastoma organoids and evaluated their viability via standard histology-based approaches: assessing cellular density (hereafter, cellularity) by hematoxylin staining and cell death by IHC for cleaved caspase 3 (CC3) (Figure 2B). With these assays established, we sought to test whether increased organoid topographical heterogeneity is observed in organoids following treatment with ionizing radiation (RT) and temozolomide (TMZ). This combination represents the standard-of-care therapy for gliomas and has been previously shown to exert antitumor activity in glioblastoma organoids, as measured by conventional methods that require tissue fixation^7^. To use Apex Imaging data to construct a visual representation of the organoid surface and quantify changes in “roughness” following chemoradiotherapy, we constructed an algorithm called SSDquant. Applying this computational tool to Apex Imaging results from untreated and treated organoids revealed striking differences in their topographical variability. The latter showed marked perturbations to the organoid surface, which aligned with visual observations of decreased organoid structural integrity following combination therapy.

To test the correlation between topographical heterogeneity and gold standard histology-based assays of tissue viability, we employed a glioma organoid model that was derived from an WHO grade 3 oligodendroglioma specimen. To generate a series of glioma organoids with wide variance in viability profiles, we treated 14 organoids with RT, the poly-ADP-ribose polymerase inhibitor Olaparib, or both together. Olaparib is currently being investigated in a number of ongoing clinical trials for glioma therapy. We then performed Apex Imaging on this suite of organoids and immediately fixed them for histological analysis. As expected, we observed a range of cellularity and cell death rates by CC3 IHC, as depicted in representative organoids (Figure 2E). Comparing these assays with natural logarithm-transformed SSD values revealed statistically significant negative and positive correlations with cellularity and cell death rate metrics, respectively (Figures 2F and G). Intriguingly, tandem Apex Imaging and computational analysis via the SSDquant algorithm appears to capture both cell loss and cell death events in organoids, which are only individually reflected in cellularity and caspase 3 cleavage assays, respectively.

## Discussion

We present a novel imaging methodology, Apex Imaging, and a companion computational tool, SSDquant, for non-invasive surveillance of patient-derived cancer organoid viability. This approach enables high-throughput analysis of laser scanning confocal microscopy data and can be scaled to assess treatment response in a cohort of treated organoids over time. Future applications of this work may allow for rapid, semi-automated, and parallel analyses of drug libraries in large collections of patient-derived organoids. Therefore, our methodology highlights new opportunities to translate recent advances in preclinical organoid modeling into tractable strategies for personalized cancer treatment.

## Acknowledgements

The authors would like to thank Melissa Logies for her illustration.

